# Practical unidentifiability of a simple vector-borne disease model: implications for parameter estimation and intervention assessment

**DOI:** 10.1101/164079

**Authors:** Yu-Han Kao, Marisa C. Eisenberg

## Abstract

**Background:** Mathematical modeling has an extensive history in vector-borne disease epidemiology, and is increasingly used for prediction, intervention design, and understanding mechanisms. Many of these studies rely on parameter estimation to link models and data, and to tailor predictions and counterfactuals to specific settings. However, few studies have formally evaluated whether vector-borne disease models can properly estimate the parameters of interest given the constraints of a particular dataset.

**Methodology/Principle Findings:** Identifiability methods allow us to examine whether model parameters can be estimated uniquely—a lack of consideration of such issues can result in misleading or incorrect parameter estimates and model predictions. Here, we evaluate both structural (theoretical) and practical identifiability of a commonly used compartmental model of mosquitoborne disease, using 2010 dengue epidemic in Taiwan as a case study. We show that while the model is structurally identifiable, it is practically unidentifiable under a range of human and mosquito time series measurement scenarios. In particular, the transmission parameters form a practically identifiable combination and thus cannot be estimated separately, which can lead to incorrect predictions of the effects of interventions. However, in spite of unidentifiability of the individual parameters, the basic reproduction number was successfully estimated across the unidentifiable parameter ranges. These identifiability issues can be resolved by directly measuring several additional human and mosquito life-cycle parameters both experimentally and in the field.

**Conclusions:** While we only consider the simplest case for the model, without explicit environmental drivers, we show that a commonly used model of vector-borne disease is unidentifiable from human and mosquito incidence data, making it difficult or impossible to estimate parameters or assess intervention strategies. This work illustrates the importance of examining identifiability when linking models with data to make predictions, and particularly highlights the importance of combining experimental, field, and case data if we are to successfully estimate epidemiological and ecological parameters using models.

**Author Summary:** Mathematical models have seen increasing use in understanding transmission processes, developing interventions, and predicting disease incidence and prevalence. Vector-borne diseases in particular present both a challenge and an opportunity for modeling, due to the complex interactions between host and vector species. A key step in many of these studies is connecting transmission models with data to infer parameters and make useful predictions, which requires careful consideration of identifiability and uncertainty of the model parameters. Whether due to intrinsic limitations of the model structure, or practical limitations of the data collected, is common that many different parameter values may yield the same or very similar fits to the data, making it impossible to successfully estimate the parameters. This issue of parameter unidentifiability can have broad implications for our ability to draw conclusions from mechanistic models—in some cases making it difficult or impossible to generate specific predictions, forecasts, or parameter estimates from a given model and data. Here, we evaluate these questions for a commonly-used model of vectorborne disease, examining how parameter uncertainty and unidentifiability can affect intervention predictions, estimation of the basic reproduction number, and other public health conclusions drawn from the model.

## 1 Introduction

Arboviral diseases are a global threat of increasing importance. Particularly for diseases propagated by *Aedes* mosquitoes, such as dengue, chikungunya, and Zika [1, 2], incidences have been increasing at alarming rates worldwide, with over approximately 3.9 billion individuals believed to be at risk for dengue infection alone [3–5]. These increases are primarily attributed to the habitat expansion of *Aedes spp*. caused by changes in anthropogenic land use and human movement [6–11]. Given the ecology and life-cycle of *Aedes* mosquitoes, the transmission dynamics of these mosquito-borne diseases are heavily driven by complicated interactions between environmental factors [12–17]. These factors, combined with human behavior and transmission dynamics, make vector-borne diseases highly complex—presenting both challenges and opportunities for mathematical modeling [18–20]. Modeling has increasingly been viewed as a useful tool to quantify these complex transmission systems by integrating various data sources and specifying nonlinear mechanistic relationships and feedbacks. Numerous recent efforts at combating mosquito-borne diseases have directly incorporated the use of mathematical models, such as in planning for Zika and chikungunya response [21–26], and evaluation of potential vaccine candidates [27–30].

Indeed, mathematical modeling has a long history in vector-borne diseases, beginning with the original development of the Ross-Macdonald or so-called Susceptible-Infectious-Recovered (SIR) model to examine malaria [31], and expanding to account for an enormous range of factors affecting both human and vector population dynamics [32,33]. A wide range of modeling approaches, including ordinary and partial differential equations (ODE and PDE) [34, 35] as well as agent/individual-based models have also been applied to these questions [27, 36–39]. Common goals for many of these modeling efforts have been to make quantitative predictions of disease dynamics and to estimate the underlying mechanistic parameters [26, 40–42].

To do so often requires using parameter estimation to connect these models with disease data, mainly using incidence or prevalence over time in humans. An important step in this process is examining parameter identifiability, the study of whether a set of parameters can be uniquely estimated and what parameter information may be gleaned from a given model and data set. Unfortunately, under many circumstances, the underlying model parameters are unidentifiable (also denoted non-identifiable), so that many different sets of parameter values produce the same model fit. The unidentifiability (non-identifiability) may be due to the model structure (i.e. structural non-identifiability) or the constraints of a specific dataset (i.e. practical unidentifiability). In either case, the data does not provide sufficient information for unique parameter estimation. Incorrect parameter estimates and ignorance of the uncertainty in prediction from an unidentifiable model can result in misleading epidemiological inferences, which could further lead to failures of public health interventions.

However, in spite of the abundance of transmission models in mosquito-borne diseases and the common use of parameter estimation in fitting these models to data, relatively few efforts have been made to examine questions of parameter identifiability in these models. [43–50]. Two studies that directly evaluated the issues include: Denis-Vidal, Verdière, and colleagues assessed the structural (theoretical) identifiability of a chikungunya transmission model assuming all the states in human population and mosquito larva are observable [46, 49]; Tuncer et al. [50] examined both structural and practical identifiability of a within-to-between host model of Rift Valley fever, addressing how the multi-scale nature of such immuno-epidemiological problems affects model identifiability.

Building on these results, we examine the identifiability of a simple compartmental model based on the Ross-Macdonald framework with various scenarios of measurement assumption [51]. This model is commonly used for both theoretical [52–55] and applied epidemiological studies in a wide range of settings [56–61], and is often used in an expanded form where temperature or environmental dependence is explicitly included [62–65]. We consider the structural and practical identifiability of this model in the baseline case without explicit environmental drivers, using dengue incidence data in Kaohsiung, Taiwan as a case study. Additionally, the inclusion of mosquito population data has been considered helpful for parameter estimation in models involving mosquito life cycles [33, 63, 66, 67]. However, obtaining mosquito population data is difficult in practice: it requires substantial time and resources which are often limited; spatial and behavioral variability in mosquito populations pose significant logistic challenges as well. Therefore, we also evaluate whether and to what degree that alternative mosquito data available in the field will reduce parameter uncertainty and improve model inference on mosquito control strategies. Finally, we present an example showing the consequences of ignoring unidentifiability in model-based intervention design.

Vector-borne disease modeling is often complex, and has been widely used in forecasting and the design of interventions [26, 28, 68–71]. Through our simple model, we hope to draw attention to identifiability issues in vector-borne disease models and their implications in the application of models with more complexity.

## 2 Methods

In the following sections, we will describe the model development, identifiability analysis, and parameter estimation processes. The flow chart in Fig. 1 summarizes the overall analytical process. The model and analyses were implemented in Python 2.7.10, with code available at https://github.com/epimath/dengue_model.

**Fig 1.**
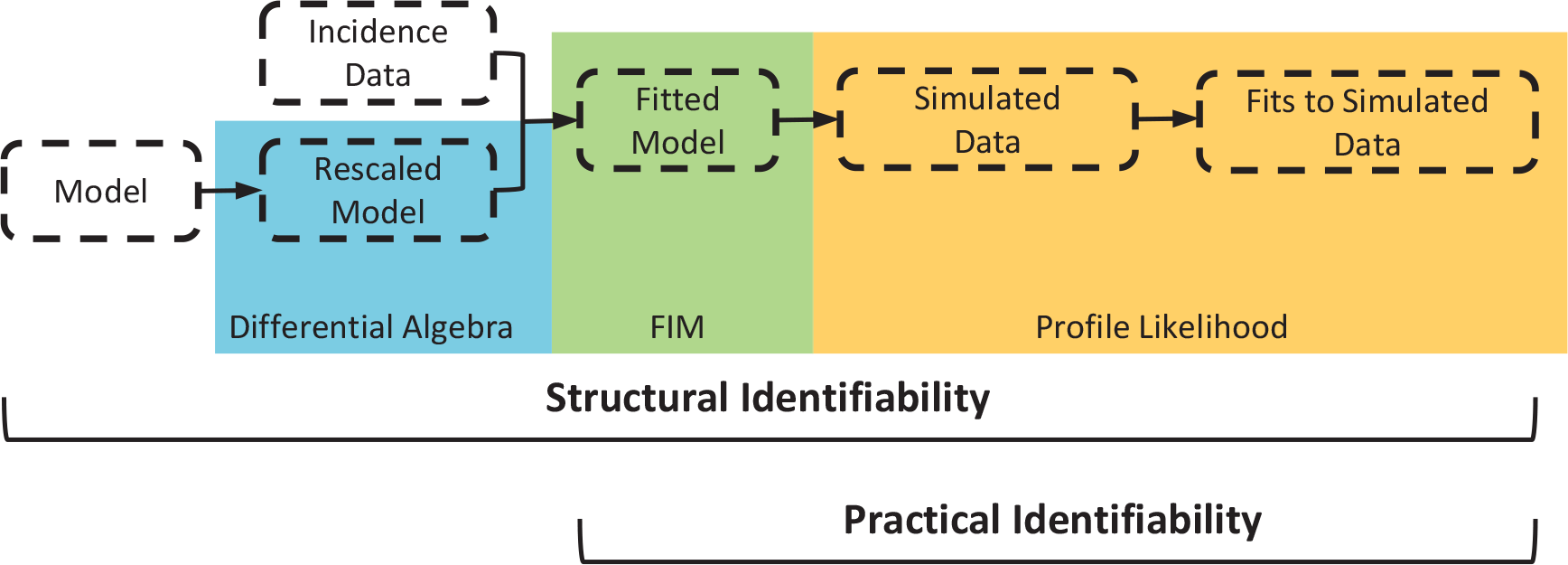
Summary of the parameter estimation and identifiability analysis process.

### 2.1 Model

Our SEIR-based model is adapted from [51, 63, 72], and shown in Fig 2. We chose this model mainly because of its simplicity as well as its potential to be used for intervention design and epidemic prediction accounting for environment factors [59–61, 63–65, 68, 72]. The model includes the disease transmission process between the human (*h*) and mosquito (*m*) populations. In addition, we specify an aquatic stage of mosquitoes combining larvae and pupae (*A*). These larvae/pupae then grow into adults (*S_m_
*) and leave the aquatic environment. Since dengue virus is transmitted by the female, we only consider female mosquitoes in the model. The susceptible adult mosquitoes become infected and enter compartment *E_m_
* by having blood meals from infectious human beings carrying the dengue virus (*I_h_
*). After the extrinsic incubation period (8-12 days) [73–75], the infected mosquitoes are capable of transmitting the virus and stay contagious during their lifetime. Susceptible human individuals (*S_h_
*) can be infected (*E_h_
*) through bites from the mosquitoes, and then become infectious (*I_h_
*) after a 4-10 day intrinsic incubation period [73–75]. With proper treatment, individuals in the infectious stage can recover from dengue and are considered immune in the model. Note that multiple serotypes are not considered in the model, so potential interactions or antibody-dependent enhancement between serotypes are not included. We assume there is only mosquito-to-human and human-to-mosquito transmission in the model given the relatively low probability of other transmission pathways [73].

**Fig 2.**
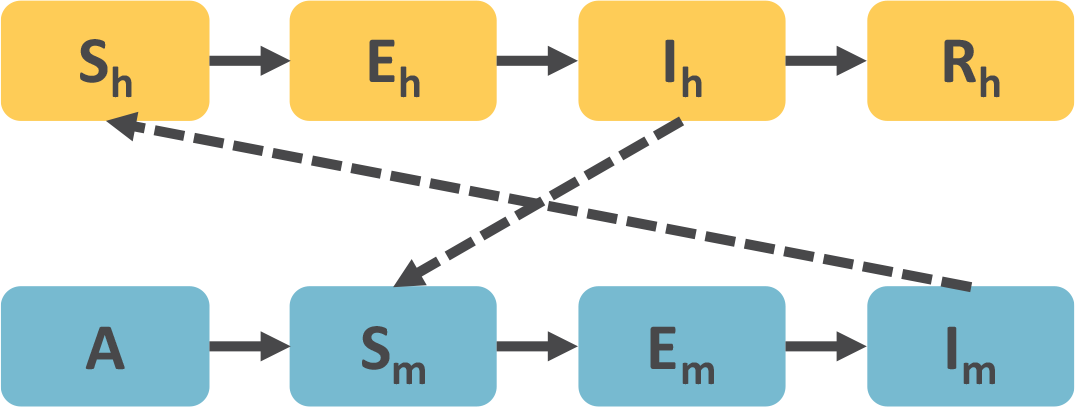
Diagram of the SEIR-based model. Subscript *h* indicates human, *v* indicates vector, and *S, E, I, R* represent susceptible, latent (exposed), infectious, and recovered humans or adult mosquitoes. *A* represents immature mosquitoes (larvae and pupae).

#### 2.1.1 Model Equations

In the model, we assume a constant human population (*N* = *S_h_
* + *E_h_
* + *I_h_
* + *R_h_
*). We also consider all variables in units of individuals (i.e. humans, mosquitoes, and pupae/larvae).

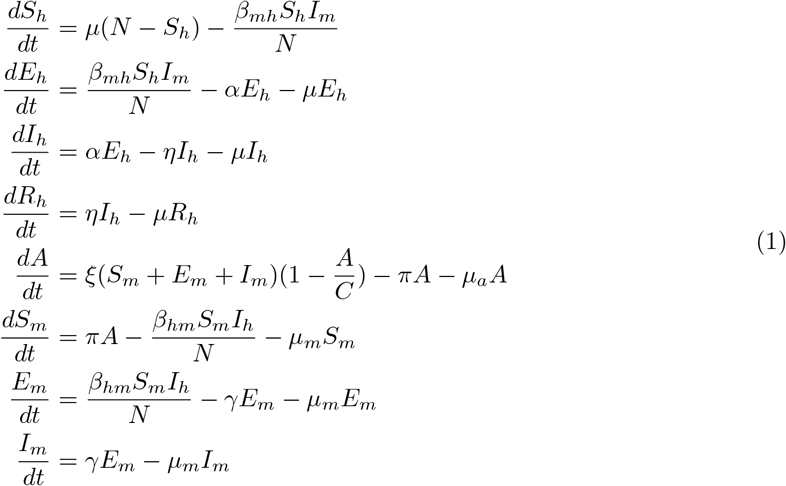

It should be noted that *β_mh_
* and *β_hm_
* are transmission rates between host and vector populations, which are the products of average bites per mosquito and the probability of successful transmission per infected mosquito bite. *C* is the carrying capacity of aquatic environment, and *π* is the maturation rate to adult mosquitoes. We also include a parameter to account for underreporting in human incidence and prevalence, so that the incidence in the model is measured as *y_h_
* = *κ_h_αE_h_
*, where *κ_h_
* is the reporting fraction. Similarly, for counts and prevalence of mosquitoes, we assume that only a small fraction of the total mosquitoes are counted, assumed to be *κ_a_
* and *κ_m_
* for aquatic immature and mature mosquitoes, respectively. This yields the (simulated) observed immature mosquitoes tobe *y_a_
* = *κ_a_A* and observed adult mosquitoes to be *y_m_
* = *κ_m_
*(*S_m_
* + *E_m_
* + *I_m_
*). Descriptions of the other parameters are given in Table 1.

**Table 1.**
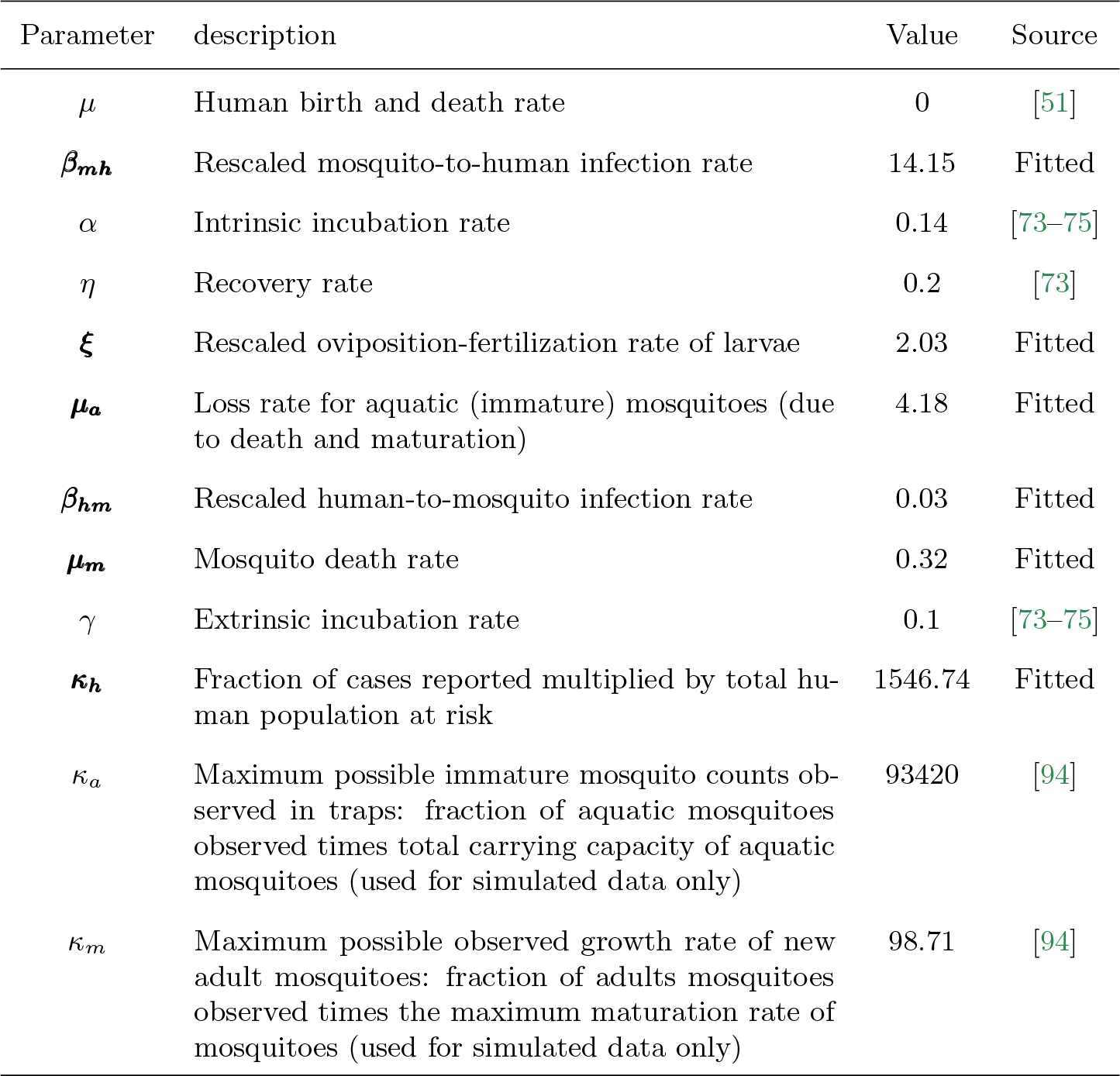
Parameter estimates and values. Estimated parameters are marked in bold; confidence bounds and uncertainty for the fitted parameters are examined further below.

| Parameter | description | Value | Source |
| --- | --- | --- | --- |
| $\mu$ | Human birth and death rate | 0 | [51] |
| **$\beta_{mh}$** | Rescaled mosquito-to-human infection rate | 14.15 | Fitted |
| $\alpha$ | Intrinsic incubation rate | 0.14 | [73–75] |
| $\eta$ | Recovery rate | 0.2 | [73] |
| **$\xi$** | Rescaled oviposition-fertilization rate of larvae | 2.03 | Fitted |
| **$\mu_a$** | Loss rate for aquatic (immature) mosquitoes (due to death and maturation) | 4.18 | Fitted |
| **$\beta_{hm}$** | Rescaled human-to-mosquito infection rate | 0.03 | Fitted |
| **$\mu_m$** | Mosquito death rate | 0.32 | Fitted |
| $\gamma$ | Extrinsic incubation rate | 0.1 | [73–75] |
| **$\kappa_h$** | Fraction of cases reported multiplied by total human population at risk | 1546.74 | Fitted |
| $\kappa_a$ | Maximum possible immature mosquito counts observed in traps: fraction of aquatic mosquitoes observed times total carrying capacity of aquatic mosquitoes (used for simulated data only) | 93420 | [94] |
| $\kappa_m$ | Maximum possible observed growth rate of new adult mosquitoes: fraction of adults mosquitoes observed times the maximum maturation rate of mosquitoes (used for simulated data only) | 98.71 | [94] |

#### 2.1.2 Rescaled Model

Transmission models such as the one considered here can often be rescaled without changing the observed output. For example, in this model we could rescale the human variables to be larger (thereby also increasing the population size *N*), but reduce the reporting rate (*κ_h_
*) and adjust the value of *β_mh_
* to yield the same apparent observed number of cases over time from the model. However, because each of these parameters (the reporting rate, transmission parameters, and size of the total population at risk) are all unknown parameters for our model, there is an inherent (structural) unidentifiability of these parameters, so that they cannot all be estimated simultaneously (i.e. for any population size, we can set *β_mh_
* and the reporting rate to yield the same observed number of cases). Similar issues can be found in the mosquito equations as well.

One way to correct these types of identifiability problems in the model is to rescale the model variables (e.g. *S_h_
*, *E_h_
*, *I_h_
*, *S_m_
*, etc.) by model parameters such as the total population size (in many cases this is equivalent to nondimensionalizing the system). In this case, we re-write the human model variables to be in terms of fraction of the population instead of numbers of individuals, e.g. letting the new variable for susceptible humans be: *S͂*
_
*h*
_ = *S_h_/N* (and similarly for *E_h_, I_h_
*, and *R_h_
*). We also normalize the larvae *A* by their carrying capacity *C* (letting *Ã* = *A/C*) and the remaining variables (*S_m_, E_m_
*, and *I_m_
*) by both *C* and *π* (i.e. letting *S͂*
_
*m*
_ = *S_m_/*(*Cπ*)). Rewriting the equations and omitting the *∼*’s yields:

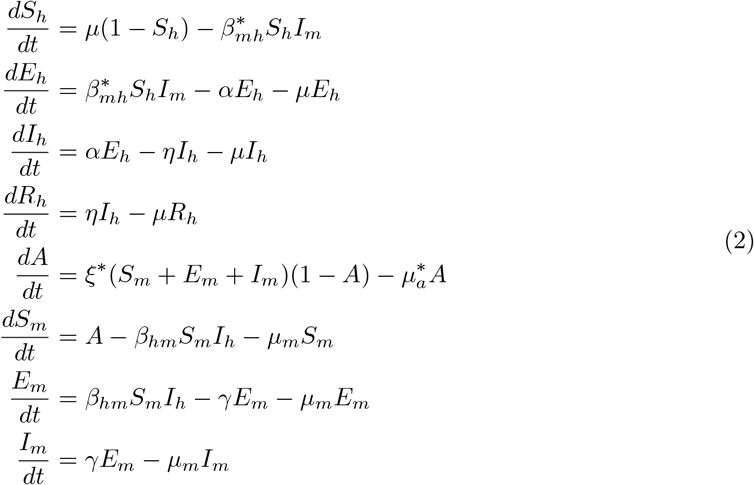

where 
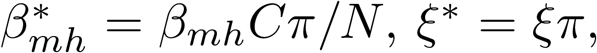
 and 
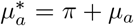
. Similarly, the reporting rate parameters are now *

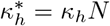

*, *

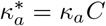

*, and *

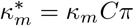

*, so that the observed human cases or mosquito counts are the same as in the original model. Doing so allows us to reduce the number of parameters explicitly included in the model and correct some of the immediately apparent identifiability issues. We will show in Section 2.3 below that this also resolves the overall structural identifiability of the model.

For the rescaled model, we can calculate the disease-free steady state value for the mosquito population, with nonzero solution
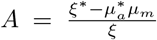
 yielding 
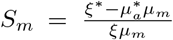
 at the disease-free equilibrium. We assume steady state for the initial condition of mosquito population. For simplicity, from here on we will work entirely with the rescaled model (Eq. (2)), and so omit the *∗*’s on the rescaled parameters in the subsequent sections.

#### 2.1.3 Basic Reproduction Number

The basic reproduction number (ℛ_0_) is the total number of secondary cases generated by introducing a single infected individual into a completely susceptible population [76, 77]. Mathematically, ℛ_0_ is a threshold parameter controlling the stability of the disease-free equilibrium given by an entirely susceptible human and mosquito population. Using the next generation matrix [76], we construct

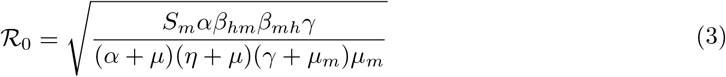

### 2.2 Parameter Estimation

#### 2.2.1 Data

Weekly incidence of dengue cases since 1998 is available from the Taiwan National Infectious Disease Statistics System of Taiwan Centers for Disease Control (CDC) [78]. Confirmed dengue cases are reported from local hospitals and are released every week to the CDC online platform. In the study, we used 2010 dengue incidence data in Kaohsiung, the main city in southern Taiwan. Dengue outbreaks in Taiwan always start from and are often confined to the south because of the favorable environment for *Aedes spp*. Kaohsiung is usually the main epidemic area during outbreaks, and also has annual outbreaks regularly [79]. The 2010 epidemic curve of dengue in Kaohsiung is very typical with one main peak. Since our model does not handle spatial heterogeneity and multiple strains, we chose to focus only on the 2010 data in Kaohsiung for these analyses.

#### 2.2.2 Parameter Estimation

We neglect population birth/death dynamics in the model (*µ* = 0) because the outbreak only lasts for 32 weeks. We also fix *α* and *γ* as 0.14 and 0.1 respectively based on previous studies [73–75], and let *η* be 0.1 since the infection usually lasts for about 10 days [73]. We estimated the remaining 6 parameters using weekly dengue incidence in Kaohsiung with least squares, assuming normally distributed errors. Nelder-Mead from NumPy in Python 2.7.10 was used for the estimation process.

#### 2.2.3 Simulated Data

As discussed in Identifiability Analysis below, we also simulated noise-free data using the fitted model from previous step. These data were generated by simulating the given variables at either daily or weekly frequency. This allowed us to examine identifiability of the model in a case where the “true” parameters are known (so that errors in estimation can be assessed) and to consider a range of alternative measurement scenarios examining how adding different types of mosquito count data might improve parameter identifiability. We synthesized the following four alternative simulated data sets corresponding to different surveillance methods available in the field––dengue incidence, ovitrap/house index, BG-trap, and Gravid trap, respectively:
- Scenario 1: human incidence data only, given by *y_h_
* = *κ_h_
*(*αE_h_
*) (integrated to a weekly cumulative incidence)
- Scenario 2: human incidence data (*y_h_
*) and daily aquatic (immature) mosquito counts, given by *y_a_
* = *κ_a_A*
- Scenario 3: human incidence data (*y_h_
*), aquatic mosquito counts (*y_a_
*), and daily adult mosquito counts, given by *y_m_
* = *κ_m_
*(*S_m_
* + *E_m_
* + *I_m_
*)
- Scenario 4: human incidence data (*y_h_
*), aquatic mosquito counts (*y_a_
*), and daily adult mosquito counts broken down by infection status, allowing us to break *y_m_
* into *y_ms_
* = *κ_m_S_m_
* and *y_mei_
* = *κ_m_
*(*E_m_
* + *I_m_
*).

#### 2.2.4 Estimation with Simulated Data

For parameter estimation using the simulated data, we fit the model with weighted least squares to account for the different scales for mosquito and human data sets. The weights are the same for each point within each individual dataset (i.e. weighted by the average data value).

### 2.3 Identifiability Analysis

We evaluated the structural and practical identifiability of the parameters, given the model and different possible data sets described above. We will give a brief overview of the identifiability definitions and methods used here. For a more complete review, please refer to [80–83].

In general there are two types of identifiability: *structural identifiability*, which examines the best-case scenario of perfectly measured, noise-free data, in order to reveal the inherent, theoretical identifiability derived from the model structure itself; and *practical identifiability*, which examines how parameter identifiability fares when real-world data issues such as noise, sampling frequency, and bias are considered [82]. When a model is unidentifiable, model parameters usually form *identifiable combinations*, which are combinations of parameters that are identifiable even though the individual parameters in the combinations are not.

#### 2.3.1 Structural Identifiability Analysis

We first examined structural identifiability using two approaches: differential algebra [81, 84–87] and the Fisher information matrix [80, 88–90]. A short overview of both methods, formal definitions, and examples are provided in the Supporting Information.

In brief, the differential algebra approach is an analytical method which examines whether is possible, from the model equations and variables measured, to uniquely determine (estimate) the parameter values. The approach is based only on the model and data structure—it assumes perfect, noise-free data, without consideration of real-world issues of noise, bias, or sampling. This represents an idealized, best-case scenario; however many biological and epidemiological models are structurally unidentifiable, making this a useful first step in examining the parameter information available for a given model and data.

The differential algebra approach provides global results of model structural identifiability and closed forms of the relationships between parameters, but it is usually very computationally expensive. The Fisher information matrix (FIM) can be used as numerical or analytical approximation to examine structural identifiability for a single point in parameter space (local results), for example, by using very finely sampled simulated data, as discussed in more detail in [90, 91]. Given that the FIM is often used as a numerical rather than analytical method, there can be limited generalizability across the parameter space. However, it is significantly faster and less computationally intensive than differential algebra approach.

Here, we test the four simulated data scenarios given above, using the differential algebra approach when possible (using both Mathematica code as well as the freely available packages COMBOS [92] and Daisy [93]), and the FIM when the differential algebra approach was too computationally intensive to converge to a solution.

#### 2.3.2 Structural and Practical Identifiability Using the Profile Likelihood

Another way to assess identifiability is the profile likelihood [82]. Taking **p** = {*θ*
_1_, · · ·, *θ_p_
*} as the parameters to be estimated, we fix a parameter (*θ_i_
*) across a range of values, which is denoted as [*min*(*θ_i_
*), *max*(*θ_i_
*)], and fit the remaining parameters {*θ_j_
* |*j* = 1, …, *p, j* ≠ *i*} using the likelihood function ℒ for each value of *θ_i_
* in [*min*(*θ_i_
*), *max*(*θ_i_
*)]. In our case, least squares is used to compute the best-fit values of *θ_i_
*s, constituting the likelihood profile for the fixed parameter. A minimum in the profile likelihood indicates structural identifiability. A parameter is structurally unidentifiable when its likelihood profile is flat and is practically unidentifiable when the curvature of its likelihood profile is shallow [82, 90]. However, the degree of shallowness for a profile is a gradated question, so there is often some question of where to set a threshold for practical unidentifiability. In order to better decide whether the profile is “flat”, we construct an approximate 95% upper confidence bound for the profile likelihood 
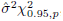
 where 
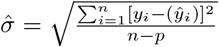
 with *n* denoting the number of observations, *p* the number of parameters to be estimated, and *y* and ŷ the observations and model trajectory respectively [82]. Using profile likelihood method, we examine the identifiability of the model with the four simulated data scenarios as well as the real dengue case data from 2010 in Kaohsiung, Taiwan.

## 3 Results

### 3.1 Model Fitting and Parameter Estimation

Using 2010 dengue incidence data in Kaohsiung, the fitted model was able to describe the general trend of the dengue epidemic. The left panel in Fig. 3 shows the dengue incidence data in 2010 and the fitted epidemic curve (*y_h_
*). The model captures the overall epidemic size and the long tail at the end (though it overshoots for some of the tail). The fitted parameter values are given in Table 1.

**Fig 3.**
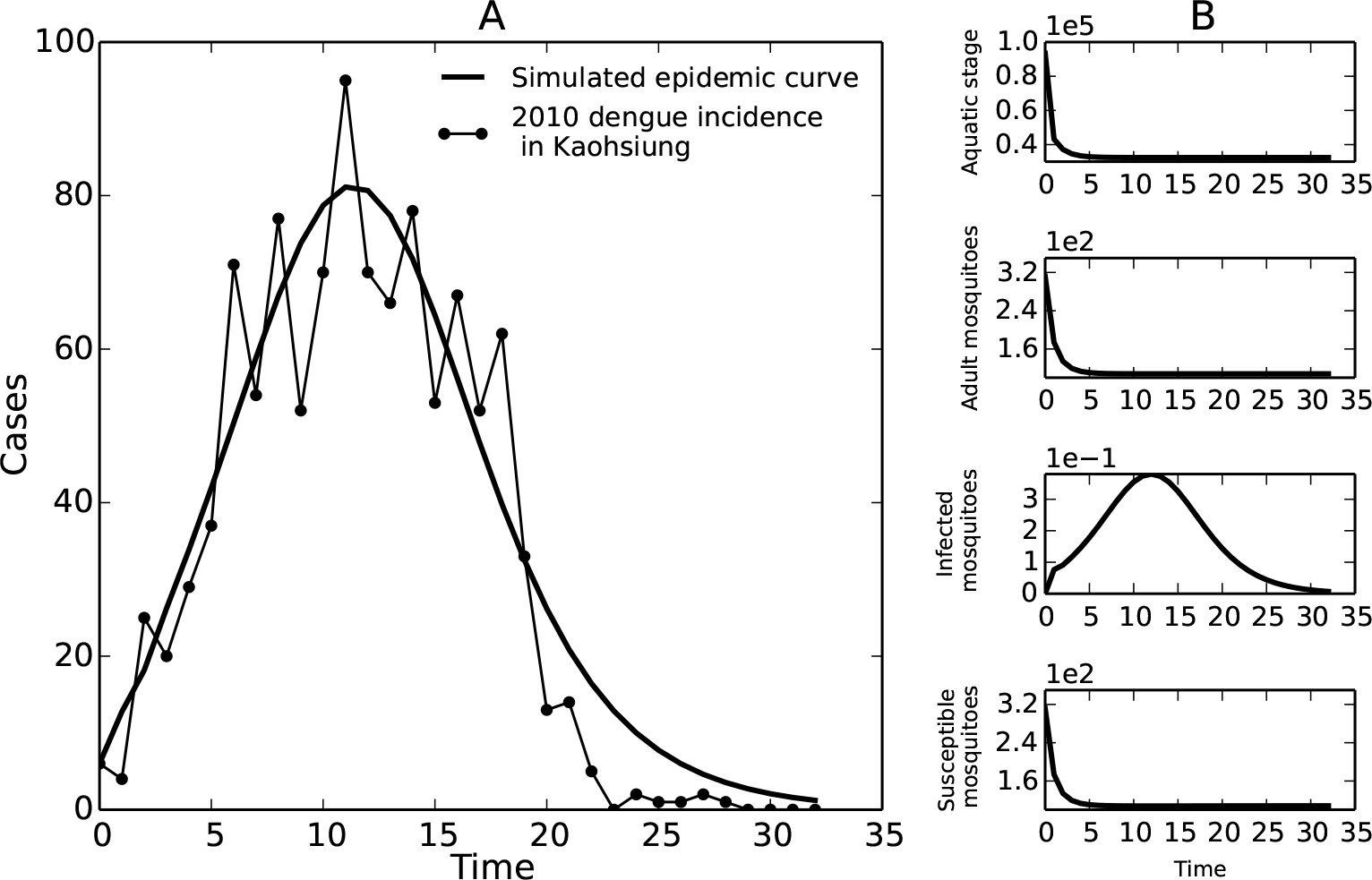
(A) model (dotted line) fitted to weekly incidence data (black circles) in Kaohsiung, Taiwan (2010); (B) simulated mosquito population data corresponding to Scenarios 2-4.

As described in the methods, we also simulated both human and mosquito population data which is potentially collectible in the field. The simulated mosquito population data included *y_a_
* (aquatic stage), *y_m_
* (adult mosquitoes), and *y_ms_
* (susceptible mosquitoes) and *y_mei_
* (infected mosquitoes), shown in Fig. 3 (right panel). The fitted model and these simulated data were used for the following identifiability analyses.

### 3.2 Differential Algebra and Fisher Information Matrix (FIM)

Using the differential algebra approach, we tested the best-case scenario including all the possible data sets from field, i.e. Scenario 4: dengue incidence, aquatic mosquito counts, infected mosquitoes, and susceptible mosquitoes. With these four types of data together, we proved that the model is structurally identifiable. The detailed proof can be found in the Supporting Information section. However, we were not able to apply the differential algebra method to the remaining three scenarios, due to computational limitations. Therefore, we constructed the FIM to examine the structural identifiability of the model with all scenarios (Scenarios 1-4), using simulated, noise-free dengue incidence and mosquito counts. The FIMs for all the scenarios were full-rank (rank=6, the number of parameters to be estimated), indicating that the model is locally structurally identifiable at the fitted values in Table 1.

### 3.3 Profile Likelihood of Estimated Parameters

The parameter profile likelihoods for both the dengue incidence data in 2010, Kaohsiung and the noise-free, simulated incidence data were very similar, with the Scenario 1 profiles shown in Fig. 4 and the Kaohsiung data in Supporting Information S4 Fig. Taking *β_mh_
* in Fig. 4 as an example, the star represents the weighted sum of squared error (SSE) of the original fitted parameter values, and the dots are the SSE after adjusting the *β_mh_
* value and re-fitting the rest of the parameters. The dashed lines are the thresholds for the approximate 95% confidence bound of the profile likelihood. In principle, the profile likelihood curves of identifiable parameters should cross the thresholds on either side of the minimum (star), and the parameter values where they cross would be the confidence bounds. In this case, all the profiles are flat, meaning the fits are very similar regardless of the changing parameter values, and the confidence bounds are effectively infinite in one or both directions. This result would initially appear at odds with the structural identifiability of the model we showed earlier; however, upon zooming in the profiles, we can see there are minima in each profile (Supporting Information S5 Fig). This suggests that although the model is structurally identifiable (consistent with the results from differential algebra and FIM approaches), it is not practically identifiable. To investigate the sources of this practical unidentifiability, we generated scatter plots of each pair of parameters, to evaluate whether any parameters are related to one another and form practically identifiable combinations. We were particularly interested in the pair *β_mh_
* and *β_hm_
*—since they form a product in ℛ_0_, they could potentially compensate for one another and maintain the same overall magnitude of the epidemic. Indeed, these two parameters do appear to follow an approximate product relationship in their profiles, as illustrated in Fig. 5. In addition, there was a strong linear relationship between *ξ* and *µ_a_
*, which are the parameters controlling the size of aquatic mosquito population. The remaining parameter relationships are shown in Supporting Information.

**Fig 4.**
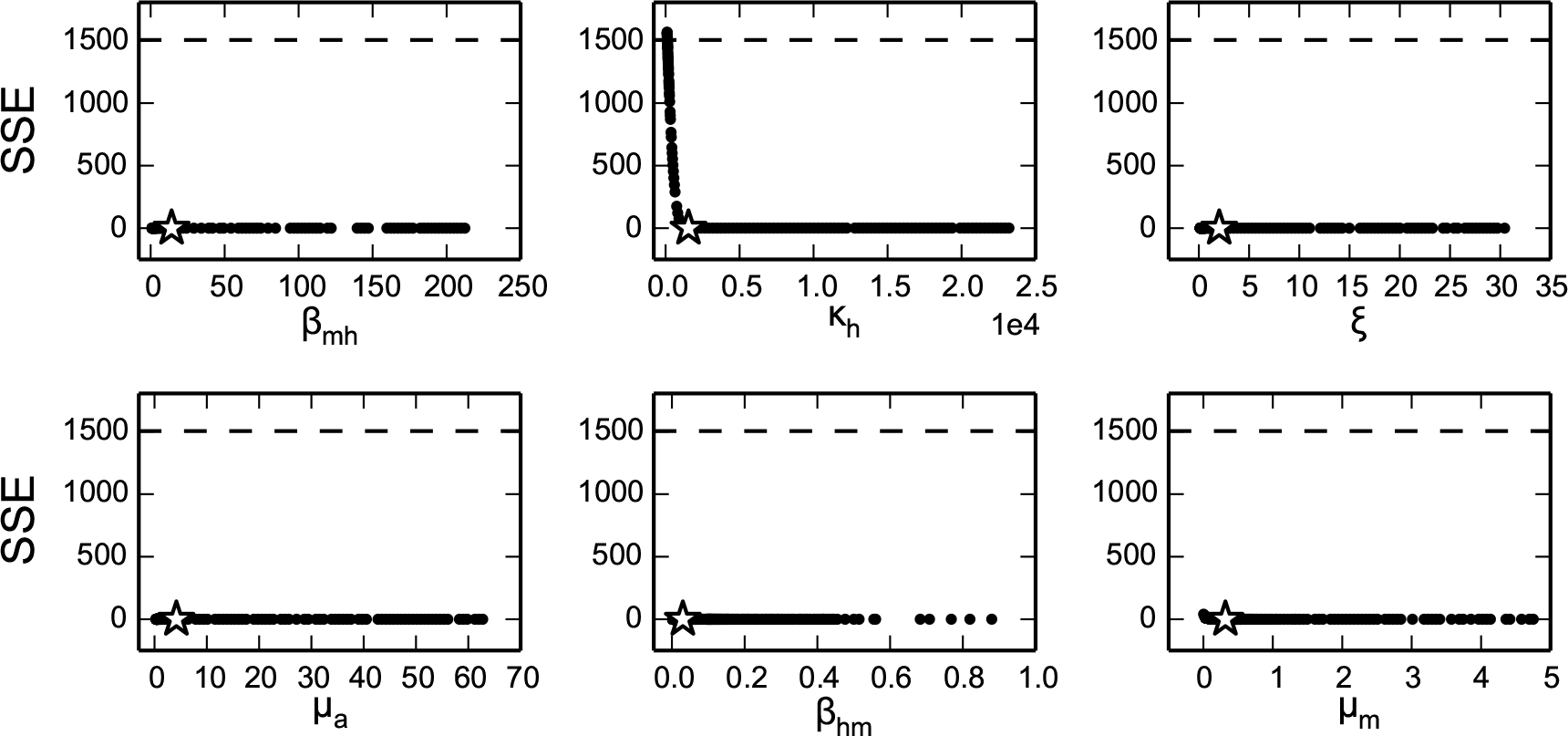
Profile likelihoods (black circles) assuming simulated, noise-free human incidence data (Scenario 1). Stars indicate the minimum sum of squared error (SSE) and dashed lines indicate the threshold for 95% confidence bounds. All six fitted model parameters are practically unidentifiable, with shallow minima which do not cross the confidence interval threshold within realistic biological ranges (zoomed in versions of the profiles showing the minima are given in the Supplementary Information).

**Fig 5.**
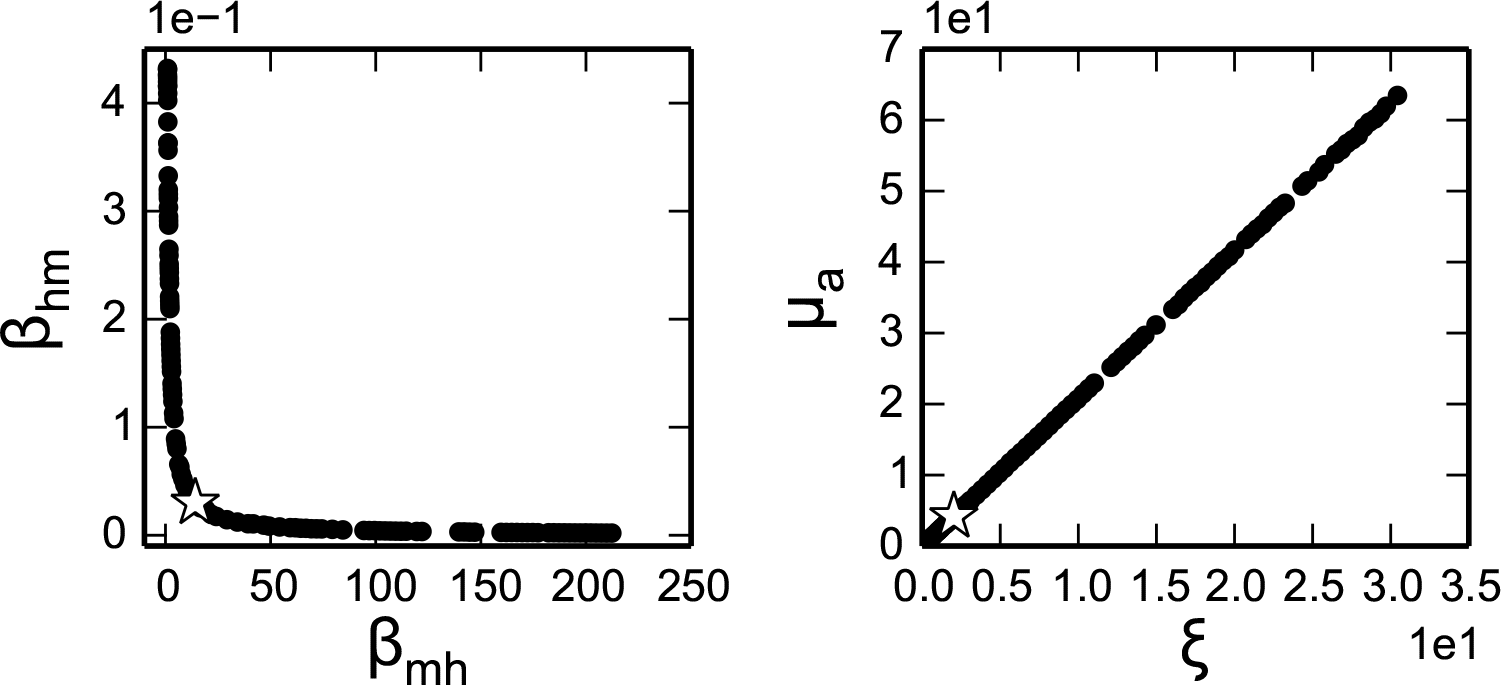
Parameter relationship scatter plots derived from the Scenario 1 profiles shown in Fig. 4, showing the relationships between *β_hm_
* and *β_mh_
* as *β_mh_
* is profiled and between *ξ* and *µ_a_
* as *ξ* is profiled. The two parameters in each pair compensate for one another, leading to the flat profile observed in Figure 4.

### 3.4 Profile Likelihood with Simulated Mosquito Data

To evaluate whether including mosquito data collection could enhance model identifiability, we computed profile likelihood of the parameters using simulated mosquito population data sets (Scenarios 2, 3 and 4). A zoomed-in comparison between the *β_mh_
* profiles of Scenario 1 (only human incidence data), Scenario 2 (adding larva data), Scenario 3 (adding larva and adult mosquito data), and Scenario 4 (adding larva, adult mosquito and infected mosquito data) is shown in Fig. 6. The profile was improved after adding mosquito information, as the curve slightly tilts up on the right-hand side and becomes higher on the left-hand side. However, the profiles including mosquito population data still do not exceed the 95% confidence threshold within a very wide range of *β_mh_
*, implying that in practice there is not much obvious improvement on the profile likelihood after including mosquito surveillance data (Supporting Information S3 Fig). We note that the small deviations from the profile curve are due to non-convergence of the estimation algorithm for some runs. The profiles for the remaining parameters are similar and are given in (Supporting Information S3 Fig). The one exception to the overall trend of practical unidentifiability was that the reporting fraction parameter for the immature mosquitoes (*κ_a_
*) was identifiable for all Scenarios where mosquito data is measured (this parameter does not appear when only human data is used).

**Fig 6.**
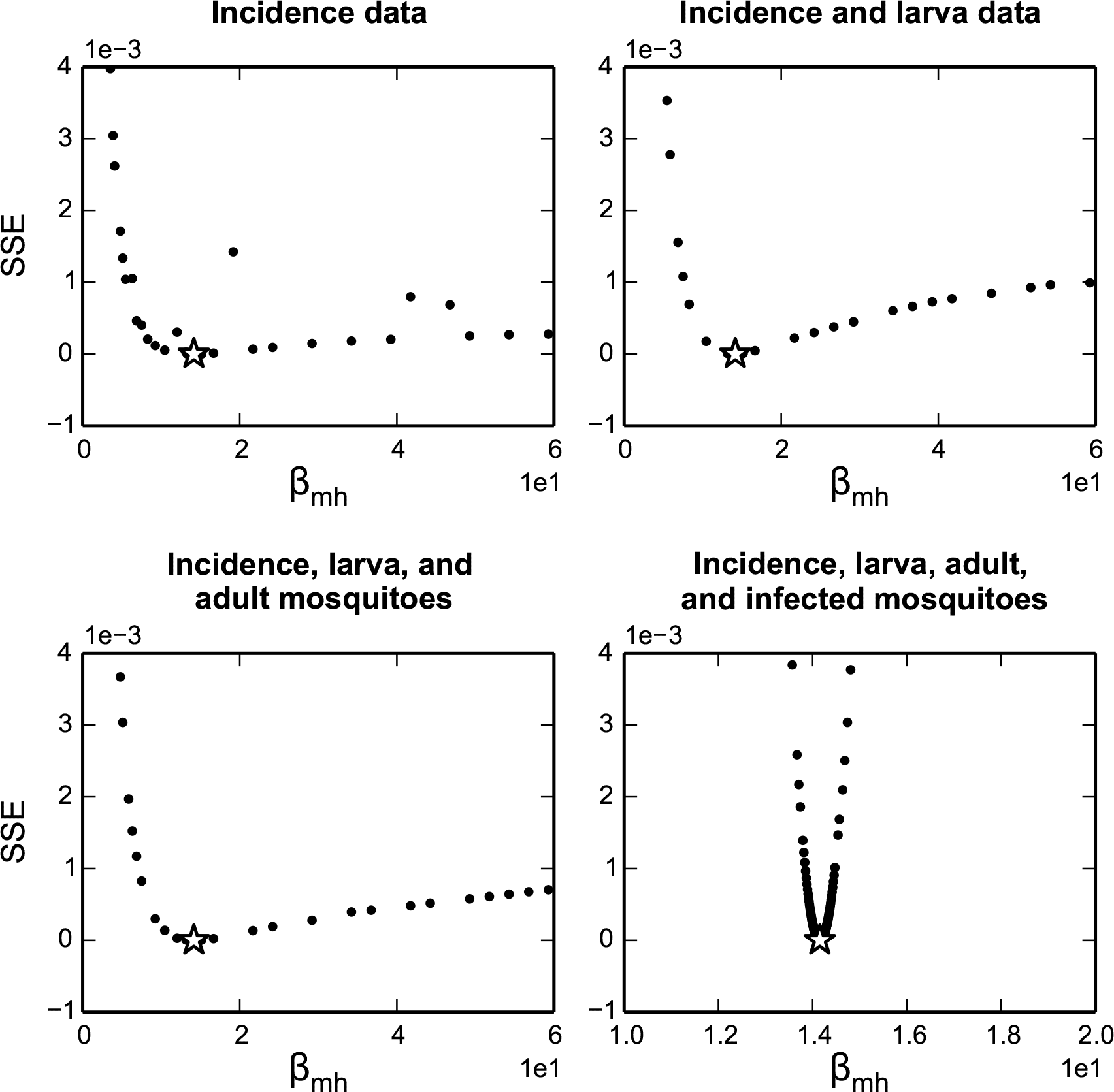
Profile likelihoods for *β_mh_
* with human incidence data only (Scenario 1), human incidence and larva count data (Scenario 2), human incidence, larva counts, and adult mosquito counts (Scenario3), and data for human incidence, larva counts, adult mosquito counts, and infected adult mosquito counts (Scenario 4).

### 3.5 Profile Likelihood with Fixed Parameters

Another way to resolve practical unidentifability is to decrease the number of parameters to be estimated, which can be done in the real world by having more information about specific parameters, such as using laboratory data to estimate the death rate for mosquito larvae. We examined this situation by fixing different sets of parameters to their originally fitted values (Table 1) and fitting the remaining parameters using synthesized dengue incidence data (Scenario 1). We demonstrate the results for the *β_mh_
* profile likelihood in Fig. 7. Given the relationship between *β_hm_
* and *β_mh_
*, one might expect fixing *β_hm_
* could resolve *β_mh_
*’s identifiability; nevertheless, the profiles indicate that fixing only one of the parameters appearing in ℛ_0_ (*β_hm_
* or *µ_m_
*) is not sufficient to make *β_mh_
* identifiable. Fixing any of other combinations of the parameters not shown in ℛ_0_ does not improve *β_mh_
*’s identifiability either. However, after fixing *β_hm_
* as well as either *µ_m_
* or the pair *ξ* and *µ_a_
*, we obtained profile likelihoods with clear minima, crossing the confidence interval threshold, suggesting with a better idea or prior knowledge about these parameters, we can make *β_mh_
* identifiable. Unfortunately, as shown in Supporting Information, the whole model does not become identifiable until we fix at least four out of six parameters of interest. A similar idea could also be incorporated in a Bayesian framework by adding sufficiently strong priors to some of the unidentifiable parameters, which could allow successful estimation of the parameters. We note that due to the model unidentifiability, the estimation would thus rely heavily on the priors.

**Fig 7.**
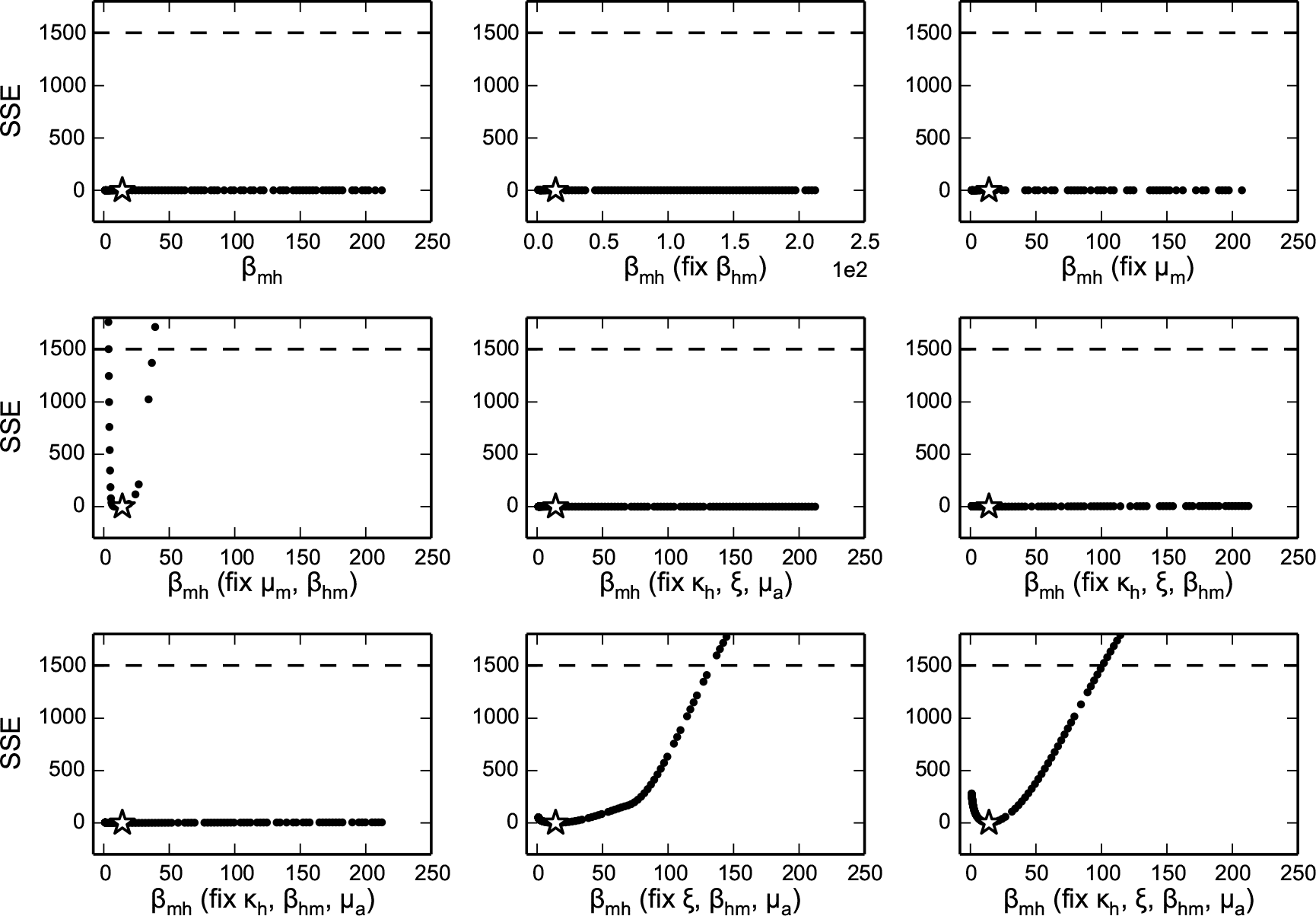
Profile likelihoods of *β_mh_
* when only subsets of *µ_a_
*, *ξ*, *κ*, *µ_m_
* and *β_mh_
* are fitted. The fixed subset (in addition to *β_mh_
*) is shown in parentheses on the x-axis.

### 3.6 Basic Reproduction Number (R_0_)

Since ℛ_0_ is an important index for understanding disease transmission and predicting future epidemics, a key question is whether we can still estimate ℛ_0_ even when the model is practically unidentifiable. As an example exploration of this question, we calculate ℛ_0_ using Eq. (3), while profiling parameters *β_mh_
* and *β_hm_
*, using Scenario 1 (human incidence data). Fig. 8 demonstrates that ℛ_0_ stays stable across the profile of *β_mh_
* and *β_hm_
* (the plots of the relationship between ℛ_0_ and other parameters are shown in Supporting Information S6 Fig. The result indicates that we can often still obtain sensible ℛ_0_ estimates from the model with human incidence data, even though we cannot properly estimate the individual parameters.

**Fig 8.**
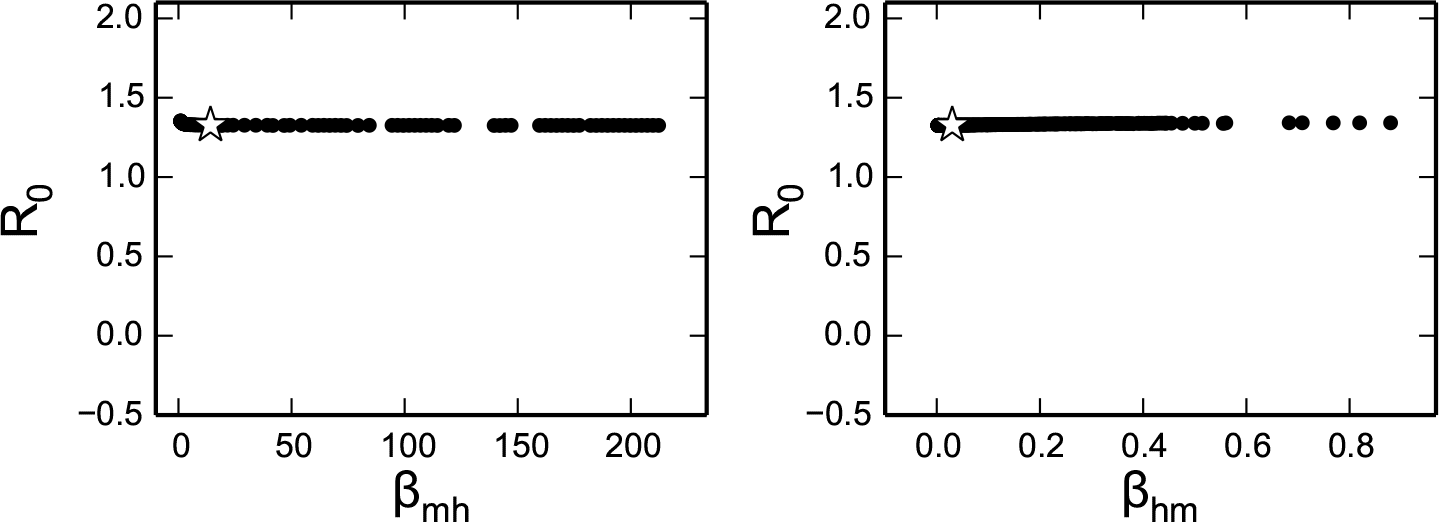
Values for ℛ_0_ as the two transmission parameters, *β_mh_
* and *β_hm_
* are varied in the profile likelihoods in Fig. 4. For each value of the profiled parameter, the plotted ℛ_0_ value uses to the best-fit values of the remaining parameters. ℛ_0_ remains relatively constant over the profiled parameter range, in spite of large changes in the parameter values.

### 3.7 Example Intervention Simulation

We implement a very naive intervention in the model to demonstrate that ignoring unidentifiability can lead to misleading outcomes. We first pick two sets of parameters from the profile in Fig. 4 that generate very similar fits (shown in Fig. 9, left panel). We then remove 10% of the aquatic (immature) mosquito population each day to simulate the population control of mosquito larvae, which is a fairly common countermeasure against dengue. With the same implementation, the responses of the two parameter sets differ substantially: one epidemic curve only decreases minimally; however, the other simulation decreases significantly and dies out at an early stage of the outbreak (Fig. 9, right panel).

**Fig 9.**
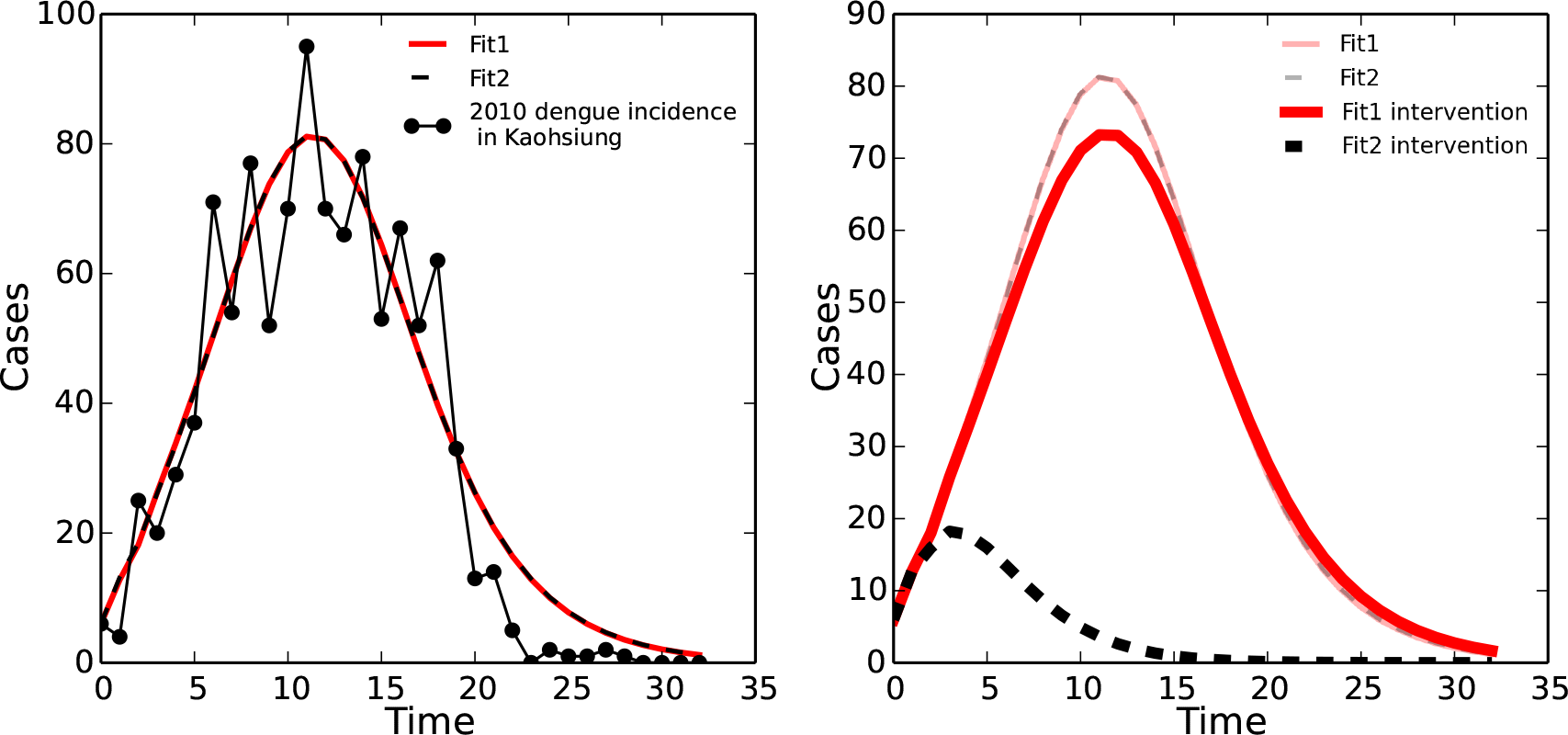
Illustration of the implications of model unidentifiability on intervention prediction. Left: two model simulations using different parameter values that give the same fit to data, based on the profiles in Fig. 4 (red solid line - original fitted parameter values from Table 1; black dashed line - [*β_mh_
* = 38.10, *κ_h_
* = 1625.42, *ξ* = 0.13, *µ_a_
* = 0.15, *β_hm_
* = 0.02, *µ_m_
* = 0.42]). Right: Simulated intervention results for both parameter sets, supposing that 10% of the aquatic (immature) mosquito population is removed at each time step.

## 4 Discussion

In this study, we explored both structural and practical identifiability of a commonly used SEIR-based model of vector-borne disease. We demonstrated that even when the model is structurally identifiable, it is likely to be difficult or impossible to estimate both human and mosquito parameters from commonly available human incidence data in a single epidemic. In other words, although the likelihood surface of the model has a single optimum, it cannot practically be distinguished from a wide range of neighboring points on the likelihood surface. Moreover, even in cases when human incidence data is combined with the types of mosquito data collected in the field, the practical identifiability of the parameters did not significantly improve. We then showed that more in-depth study of mosquito ecology and behaviors, which can give us direct information about individual parameters, was more efficient in terms of improving model identifiability. Unfortunately, obtaining accurate measurements for any of these parameters can be very difficult in practice, as they often vary depending on environmental and ecological factors such as temperature, weather events such as storms, and predation by other species [17, 94–97]. We would also need additional information on most of the parameters to make the model fully identifiable, which may not be feasible in real-world research.

In spite of these identifiability problems, the model generates very similar ℛ_0_ estimates across a range of profiled parameter values producing the same fit to the data. This means that estimation using the model may still be useful in characterizing the disease outbreak and spread, calculating vaccination coverage, and assessing the risk of vector-borne disease, even if the individual parameters cannot be determined. ℛ_0_ is an important measure that can be used to evaluate potential interventions in public health. For example, we can simulate a model that implements the intervention and compare the ℛ_0_ with and without the intervention to evaluate the potential effectiveness (e.g. by examining whether ℛ_0_ becomes less than one, or the magnitude of the reduction).

Nevertheless, we cannot solely depend on ℛ_0_ since it is possible to obtain very different predicted responses with the same intervention implementation, as shown in (Fig. 9). The two alternative parameter sets shown in Fig. 9 both fit the data equally well and have similar ℛ_0_ values (1.30 and 1.33), so that we cannot distinguish which of the predicted intervention responses is more likely. The intervention simulation used here is quite simple, but represents a commonly used control strategy. The example illustrates how a lack of consideration of parameter identifiability can potentially lead to significant errors in evaluating or comparing different intervention strategies.

This model is a simple interpretation of the world, and only assumes one outbreak and single viral strain. Despite this simple structure, we still cannot properly estimate the parameters from the model. Models with more complicated designs are more likely to be unidentifiable, underscoring the importance of taking model identifiability into account before making any inferences from the model. Identifiability analysis allows us to understand what a model and data can really tell us, and can help with planning before we invest time and resources into a experimental or field study. Even if unidentifiability is inevitable, as long as we understand the behavior, uncertainty, and the limitations of the model, mathematical models can still be powerful tools to study disease transmission. In the analyses present here, we cover a set of basic (and best-case) scenarios. More comprehensive research is needed to investigate how to handle problems that are commonly encountered in field research— different types of measurement and process noise, missing data, and data resolutions—which can further obstruct parameter estimation. Nonetheless, this work shows that parameter estimation from incidence data alone is likely to be difficult or impossible, highlighting the importance of integrating parameter information directly from experimental or field data. Given that such experimentally measured parameters usually vary as a function of environmental variables such as temperature and rainfall, future work to evaluate how model identifiability changes once the dependence is incorporated into the parameters would be a highly useful next step.

## Acknowledgments

The authors would like to thank Sarah Cherng, Lisa Lau, and Rafael Meza for their helpful comments and discussion of this work. This study was supported by the National Institutes of Health (NIH) National Institute of General Medical Sciences (NIGMS) grant U01GM110712, International Peace Scholarship from Philanthropic Educational Organization, and Study Abroad Scholarship from Ministry of Education in Taiwan.

## Supporting Information

**S1 Text Background on structural identifiability using differential algebra and the Fisher information matrix.**

**S2 Text Structural identifiability proof of Scenario 4 using differential algebra.**

**S3 Fig. Profile likelihoods with simulated human surveillance and mosquito data.** Profile likelihoods (black circles) assuming simulated human case surveillance data as well as simulated data on: immature mosquito (larvae/pupae) counts (top panel), immature and adult mosquito counts (center panel), and immature and adult mosquito counts with adult mosquito infection status (bottom panel). Stars indicate the best-fit parameter value, and dashed lines indicate the threshold for the 95% confidence intervals.

**S4 Fig. Profile likelihood with real human surveillance data.** Profile likelihoods (black circles) with human case surveillance data from 2010 in Kaohsiung, Taiwan. Stars indicate the best-fit parameter value, and dashed lines indicate the threshold for the 95% confidence intervals.

**S5 Fig. Zoomed-in profile likelihoods.** Zoomed-in views of the profile likelihoods in Figure 4 and Supporting Information S3 Fig, illustrating the minima surrounding the best fit values (best-fit parameter values given as stars).

**S6 Fig. Parameter relationships for human surveillance data profile likelihoods.** Compensatory relationships between parameters as each parameter is profiled, assuming human surveil-lance data. Some parameters show pronounced and consistent relationships with one another, indicating the possibility of a practically identifiable combination, while other parameters appear to have no relationship or noisy/inconsistent relationships depending on which parameter is profiled (possibly indicating more complex combination structures between multiple parameters rather than a single pair).

**S7 Fig. Profile likelihood when fitting subsets of parameters.** Profile likelihood (black circles) when fixing 1) *µ_m_
* and *β_hm_
*; 2)*β_hm_
*, *µ_a_
*, and *ξ*; 3)*β_hm_
*, *κ_h_ µ_a_
*, and *ξ*. Stars indicate the best-fit parameter value, and dashed lines indicate the threshold for the 95% confidence intervals.

## References

1. Musso D, Cao-Lormeau VM, Gubler DJ. Zika virus: following the path of dengue and chikungunya? Lancet. 2015;386(9990):243–244. doi:10.1016/S0140-6736(15)61273-9.

2. Benelli G, Mehlhorn H. Declining malaria, rising of dengue and Zika virus: insights for mosquito vector control. Parasitol Res. 2016;115(5):1747–1754. doi:10.1007/s00436-016-4971-z.

3. Brady OJ, Gething PW, Bhatt S, Messina JP, Brownstein JS, Hoen AG, et al. Refining the Global Spatial Limits of Dengue Virus Transmission by Evidence-Based Consensus. PLoS Negl Trop Dis. 2012;6(8):e1760. doi:10.1371/journal.pntd.0001760.

4. Bhatt S, Gething PW, Brady OJ, Messina JP, Farlow AW, Moyes CL, et al. The global distribution and burden of dengue. Nature. 2013;496(7446):504–507. doi:10.1038/nature12060.

5. World Health Organization. Dengue and sever dengue; 2017. Available from: http://www.who.int/mediacentre/factsheets/fs117/en/.

6. Wilder-Smith A, Gubler DJ. Geographic Expansion of Dengue: The Impact of International Travel. Med Clin North Am. 2008;92(6):1377–1390. doi:10.1016/j.mcna.2008.07.002.

7. Guzman MG, Halstead SB, Artsob H, Buchy P, Farrar J, Gubler DJ, et al. Dengue: a continuing global threat. Nat Rev Microbiol. 2010;8(12):S7–S16. doi:10.1038/nrmicro2460.

8. Gubler DJ. Dengue, Urbanization and Globalization: The Unholy Trinity of the 21st Century. Trop Med Health. 2011;39(4SUPPLEMENT):S3–S11. doi:10.2149/tmh.2011-S05.

9. Weaver SC. Urbanization and geographic expansion of zoonotic arboviral diseases: mechanisms and potential strategies for prevention. Trends Microbiol. 2013;21(8):360–363. doi:10.1016/j.tim.2013.03.003.

10. Wilder-Smith A. Dengue infections in travellers. Paediatr Int Child Health. 2012;32(sup1):28–32. doi:10.1179/2046904712Z.00000000050.

11. Powell JR, Tabachnick WJ. History of domestication and spread of Aedes aegypti - A Review. Mem Inst Oswaldo Cruz. 2013;108(suppl 1):11–17. doi:10.1590/0074-0276130395.

12. Hales S, de Wet N, Maindonald J, Woodward A. Potential effect of population and climate changes on global distribution of dengue fever: an empirical model. Lancet. 2002;360(9336):830–834. doi:10.1016/S0140-6736(02)09964-6.

13. Hales S, Edwards SJ, Kovats RS. Impacts on health of climate extremes. Clim Chang Hum Heal Risks responses. 2003; p. 79–102.

14. Scott TW, Morrison AC. Aedes aegypti density and the risk of dengue virus transmission. In: Ecol. Asp. Appl. Genet. Modif. Mosquitoes; 2003. p. 187–206. Available from: http://books.google.com/books?hl=en{&}lr={&}id=Sir5L1Gz23EC{&}oi=fnd{&}pg=PA187{&}dq=Aedes+aegypti+density+and+the+risk+of+dengue-virus+transmission{&}ots=cdFOU-hRSW{&}sig=oR2Dxw{_}ysML63mM7Ohl-VZUlaog.

15. Chen SC, Hsieh MH. Modeling the transmission dynamics of dengue fever: Implications of temperature effects. Sci Total Environ. 2012;431:385–391. doi:10.1016/j.scitotenv.2012.05.012.

16. Brady OJ, Golding N, Pigott DM, Kraemer MUG, Messina JP, Reiner Jr RC, et al. Global temperature constraints on Aedes aegypti and Ae. albopictus persistence and competence for dengue virus transmission. Parasit Vectors. 2014;7(1):338. doi:10.1186/1756-3305-7-338.

17. Kraemer MU, Sinka ME, Duda KA, Mylne AQ, Shearer FM, Barker CM, et al. The global distribution of the arbovirus vectors Aedes aegypti and Ae. albopictus. Elife. 2015;4. doi:10.7554/eLife.08347.

18. World Health Organization. Report of the meeting of the WHO/VMI Workshop on Dengue modeling. Geneva, Switzerland: World Health Organization; 2011. Available from: http://apps.who.int/iris/bitstream/10665/70625/1/WHO{_}IVB{_}11.02{_}eng.pdf.

19. World Health Organization. Global Strategy for Dengue Prevention and Control 2012–2020; 2012. Available from: http://apps.who.int/iris/bitstream/10665/75303/1/9789241504034{_}eng.pdf.

20. Smith DL, Battle KE, Hay SI, Barker CM, Scott TW, McKenzie FE. Ross, Macdonald, and a theory for the dynamics and control of mosquito-transmitted pathogens; 2012.

21. Moulay D, Aziz-Alaoui MA, Cadivel M. The chikungunya disease: Modeling, vector and transmission global dynamics. Math Biosci. 2011;229(1):50–63. doi:10.1016/j.mbs.2010.10.008.

22. Moulay D, Aziz-Alaoui MA, Kwon HD. Optimal control of chikungunya disease: Larvae reduction, treatment and prevention. Math Biosci Eng. 2012;9(2):369–392. doi:10.3934/mbe.2012.9.369.

23. Christofferson RC, Mores CN, Wearing HJ. Bridging the Gap Between Experimental Data and Model Parameterization for Chikungunya Virus Transmission Predictions. J Infect Dis. 2016;214(suppl 5):S466–S470. doi:10.1093/infdis/jiw283.

24. Alex Perkins T, Siraj AS, Ruktanonchai CW, Kraemer MUG, Tatem AJ. Model-based projections of Zika virus infections in childbearing women in the Americas. Nat Microbiol. 2016;1(9):16126. doi:10.1038/nmicrobiol.2016.126.

25. Kucharski AJ, Funk S, Eggo RM, Mallet HP, Edmunds WJ, Nilles EJ. Transmission Dynamics of Zika Virus in Island Populations: A Modelling Analysis of the 2013–14 French Polynesia Outbreak. PLoS Negl Trop Dis. 2016;10(5):e0004726. doi:10.1371/journal.pntd.0004726.

26. Ferguson NM, Cucunuba ZM, Dorigatti I, Nedjati-Gilani GL, Donnelly CA, Basanez MG, et al. Countering the Zika epidemic in Latin America. Science (80-). 2016;353(6297):353–354. doi:10.1126/science.aag0219.

27. Chao DL, Halstead SB, Halloran ME, Longini IM. Controlling Dengue with Vaccines in Thailand. PLoS Negl Trop Dis. 2012;6(10). doi:10.1371/journal.pntd.0001876.

28. WHO-VMI Dengue Vaccine Modeling Group. Assessing the Potential of a Candidate Dengue Vaccine with Mathematical Modeling. PLoS Negl Trop Dis. 2012;6(3):e1450. doi:10.1371/journal.pntd.0001450.

29. Aguiar M, Stollenwerk N, Halstead SB. The Impact of the Newly Licensed Dengue Vaccine in Endemic Countries. PLoS Negl Trop Dis. 2016;10(12):e0005179. doi:10.1371/journal.pntd.0005179.

30. Ferguson NM, Rodríguez-Barraquer I, Dorigatti I, Mier-y Teran-Romero L, Laydon DJ, Cummings DAT. Benefits and risks of the Sanofi-Pasteur dengue vaccine: Modeling optimal deployment. Science (80-). 2016;353(6303):1033–1036. doi:10.1126/science.aaf9590.

31. Kermack WO, McKendrick aG. Contributions to the mathematical theory of epidemics. Proc R Soc London. 1927;115(772):700–721. doi:10.1098/rspa.1927.0118.

32. Andraud M, Hens N, Marais C, Beutels P. Dynamic epidemiological models for dengue transmission: a systematic review of structural approaches. PLoS One. 2012;7(11):e49085. doi:10.1371/journal.pone.0049085.

33. Reiner RC, Perkins TA, Barker CM, Niu T, Chaves LF, Ellis AM, et al. A systematic review of mathematical models of mosquito-borne pathogen transmission: 1970-2010. J R Soc Interface. 2013;10(81):20120921. doi:10.1098/rsif.2012.0921.

34. Enduri MK, Jolad S. Dynamics of Dengue with human and vector mobility. 2014;.

35. Aldila D, Götz T, Soewono E. An optimal control problem arising from a dengue disease transmission model. Math Biosci. 2013;242(1):9–16. doi:10.1016/j.mbs.2012.11.014.

36. Li J, Zou X. Modeling spatial spread of infectious diseases with a fixed latent period in a spatially continuous domain. Bull Math Biol. 2009;71(8):2048–2079. doi:10.1007/s11538-009-9457-z.

37. Isidoro C, Fachada N, Barata F, Rosa A. Agent-based model of dengue disease transmission by Aedes aegypti populations. In: Lect. Notes Comput. Sci. (including Subser. Lect. Notes Artif. Intell. Lect. Notes Bioinformatics). vol. 5777 LNAI; 2011. p. 345–352.

38. Dommar CJ, Lowe R, Robinson M, Rodó X. An agent-based model driven by tropical rainfall to understand the spatio-temporal heterogeneity of a chikungunya outbreak. Acta Trop. 2014;129(1):61–73. doi:10.1016/j.actatropica.2013.08.004.

39. Manore CA, Hickmann KS, Hyman JM, Foppa IM, Davis JK, Wesson DM, et al. A network-patch methodology for adapting agent-based models for directly transmitted disease to mosquito-borne disease. J Biol Dyn. 2015;9(1):52–72. doi:10.1080/17513758.2015.1005698.

40. Chowell G, Diaz-Dueñas P, Miller JC, Alcazar-Velazco A, Hyman JM, Fenimore PW, et al. Estimation of the reproduction number of dengue fever from spatial epidemic data. Math Biosci. 2007;208(2):571–589. doi:10.1016/j.mbs.2006.11.011.

41. Khan A, Hassan M, Imran M. Estimating the Basic Reproduction Number for Single-Strain Dengue Fever Epidemics. Infect Dis poverty. 2014;3(1):12. doi:10.1186/2049-9957-3-12.

42. Perkins A, Siraj A, Ruktanonchai WC, Kraemer M, Tatem A. Model-based projections of Zika virus infections in childbearing women in the Americas. bioRxiv. 2016;1(9):039610. doi:10.1101/039610.

43. Mendes Luz P, Torres Codeço C, Massad E, Struchiner CJ. Uncertainties Regarding Dengue Modeling in Rio de Janeiro, Brazil. Mem Inst Oswaldo Cruz. 2003;98(7):871–878. doi:10.1590/S0074-02762003000700002.

44. Laneri K, Bhadra A, Ionides EL, Bouma M, Dhiman RC, Yadav RS, et al. Forcing versus feedback: Epidemic malaria and monsoon rains in Northwest India. PLoS Comput Biol. 2010;6(9). doi:10.1371/journal.pcbi.1000898.

45. Bhadra A, Ionides EL, Laneri K, Pascual M, Bouma M, Dhiman RC. Malaria in Northwest India: Data Analysis via Partially Observed Stochastic Differential Equation Models Driven by Lévy Noise. J Am Stat Assoc. 2011;106(494):440–451. doi:10.1198/jasa.2011.ap10323.

46. Moulay D, Verdière N, Denis-Vidal L. Identifiablility of parameters in an epidemiologic model modeling the transmission of the Chikungunya; 2012. Available from: https://hal.archives-ouvertes.fr/hal-00699172.

47. Pandey A, Mubayi A, Medlock J. Comparing vector-host and SIR models for dengue transmission. Math Biosci. 2013;246(2):252–259. doi:10.1016/j.mbs.2013.10.007.

48. Reiner RC, Stoddard ST, Forshey BM, King AA, Ellis AM, Lloyd AL, et al. Time-varying, serotype-specific force of infection of dengue virus. Proc Natl Acad Sci. 2014;111(26):E2694–E2702. doi:10.1073/pnas.1314933111.

49. Zhu S, Lilianne DV, Verdière N. Identifiability study in a model describing the propagation of the chikungunya to the human population; 2015. Available from: https://hal.archives-ouvertes.fr/hal-01166654.

50. Tuncer N, Gulbudak H, Cannataro VL, Martcheva M. Structural and Practical Identifiability Issues of Immuno-Epidemiological Vector–Host Models with Application to Rift Valley Fever. Bull Math Biol. 2016;78(9):1796–1827. doi:10.1007/s11538-016-0200-2.

51. Newton EAC, Reiter P. A model of the transmission of dengue fever with an evaluation of the impact of ultra-low volume (ULV) insecticide applications on dengue epidemics. Am J Trop Med Hyg. 1992;47(6):709–720.

52. Coutinho FAB, Burattini MN, Lopez LF, Massad E. An approximate threshold condition for non-autonomous system: An application to a vector-borne infection. Math Comput Simul. 2005;70(3):149–158. doi:10.1016/j.matcom.2005.06.003.

53. Coutinhoa FAB, Burattinia MN, Lopeza LF, Massada E. Threshold Conditions for a Non-Autonomous Epidemic System Describing the Population Dynamics of Dengue. Bull Math Biol. 2006;68(8):2263–2282. doi:10.1007/s11538-006-9108-6.

54. Garba SM, Gumel AB, Abu Bakar MR. Backward bifurcations in dengue transmission dynamics. Math Biosci. 2008;215(1):11–25. doi:10.1016/j.mbs.2008.05.002.

55. Dumont Y, Chiroleu F, Domerg C. On a temporal model for the Chikungunya disease: Modeling, theory and numerics. Math Biosci. 2008;213(1):80–91. doi:10.1016/j.mbs.2008.02.008.

56. Burattini MN, Chen M, Chow A, Coutinho FAB, Goh KT, Lopez LF, et al. Modelling the control strategies against dengue in Singapore. Epidemiol Infect. 2008;136(03). doi:10.1017/S0950268807008667.

57. Yang HM, Ferreira CP. Assessing the effects of vector control on dengue transmission. Appl Math Comput. 2008;198(1):401–413. doi:10.1016/j.amc.2007.08.046.

58. Chiroleu F, Dumont Y. Vector control for the Chikungunya disease. Math Biosci Eng. 2010;7(2):313–345. doi:10.3934/mbe.2010.7.313.

59. Pinho STR, Ferreira CP, Esteva L, Barreto FR, Morato e Silva VC, Teixeira MGL. Modelling the dynamics of dengue real epidemics. Philos Trans R Soc A Math Phys Eng Sci. 2010;368(1933):5679–5693. doi:10.1098/rsta.2010.0278.

60. Poletti P, Messeri G, Ajelli M, Vallorani R, Rizzo C, Merler S. Transmission potential of chikungunya virus and control measures: The case of italy. PLoS One. 2011;6(5). doi:10.1371/journal.pone.0018860.

61. Sardar T, Sasmal SK, Chattopadhyay J. Estimating dengue type reproduction numbers for two provinces of Sri Lanka during the period 2013-14. Virulence. 2016;7(2):187–200. doi:10.1080/21505594.2015.1096470.

62. Bartley LM, Donnelly CA, Garnett GP. The seasonal pattern of dengue in endemic areas: mathematical models of mechanisms. Trans R Soc Trop Med Hyg;96(4):387–97.

63. Yang HM, Macoris MLG, Galvani KC, Andrighetti MTM, Wanderley DMV. Assessing the effects of temperature on dengue transmission. Epidemiol Infect. 2009;137(08):1179. doi:10.1017/S0950268809002052.

64. Erickson RA, Presley SM, Allen LJS, Long KR, Cox SB. A dengue model with a dynamic Aedes albopictus vector population. Ecol Modell. 2010;221(24):2899–2908. doi:10.1016/j.ecolmodel.2010.08.036.

65. McLennan-Smith TA, Mercer GN. Complex behaviour in a dengue model with a seasonally varying vector population. Math Biosci. 2014;248:22–30. doi:10.1016/j.mbs.2013.11.003.

66. Morrison AC, Ellis AM, Garcia AJ, Scott TW, Focks DA. Parameterization and Sensitivity Analysis of a Complex Simulation Model for Mosquito Population Dynamics, Dengue Transmission, and Their Control. Am J Trop Med Hyg. 2011;85(2):257–264. doi:10.4269/ajtmh.2011.10-0516.

67. Bowman LR, Runge-Ranzinger S, McCall PJ. Assessing the Relationship between Vector Indices and Dengue Transmission: A Systematic Review of the Evidence. PLoS Negl Trop Dis. 2014;8(5). doi:10.1371/journal.pntd.0002848.

68. Yakob L, Clements ACA. A Mathematical Model of Chikungunya Dynamics and Control: The Major Epidemic on Réunion Island. PLoS One. 2013;8(3). doi:10.1371/journal.pone.0057448.

69. Patz JA, Martens WJ, Focks DA, Jetten TH. Dengue fever epidemic potential as projected by general circulation models of global climate change. Environ Health Perspect. 1998;106(3):147–53. doi:10.1289/ehp.98106147.

70. Focks DA, Barrera R. Dengue transmission dynamics: assessment and implications for control. Rep Sci Work Gr Meet Dengue. 2006; p. 92–108.

71. Kearney M, Porter WP, Williams C, Ritchie S, Hoffmann AA. Integrating biophysical models and evolutionary theory to predict climatic impacts on species’ ranges: the dengue mosquito Aedes aegypti in Australia. Funct Ecol. 2009;23(3):528–538. doi:10.1111/j.1365-2435.2008.01538.x.

72. Yang HM, Macoris MLG, Galvani KC, Andrighetti MTM, Wanderley DMV. Assessing the effects of temperature on the population of Aedes aegypti, the vector of dengue. Epidemiol Infect. 2009;137(8):1188–1202. doi:10.1017/S0950268809002040.

73. World Health Organization, Research Special Programme for Diseases and Training in Tropical. Dengue: guidelines for diagnosis, treatment, prevention, and control. In: Dengue Guidel. Diagnosis, Treat. Prev. Control; 2009. p. 160.

74. Chan M, Johansson MA. The Incubation Periods of Dengue Viruses. PLoS One. 2012;7(11):1–7. doi:10.1371/journal.pone.0050972.

75. Rudolph KE, Lessler J, Moloney RM, Kmush B, Cummings DAT. Review article: Incubation periods of mosquito-borne viral infections: a systematic review; 2014.

76. van den Driessche P, Watmough J. Reproduction numbers and sub-threshold endemic equilibria for compartmental models of disease transmission. Math Biosci;180:29–48.

77. Heffernan JM, Smith RJ, Wahl LM. Perspectives on the basic reproductive ratio. J R Soc Interface. 2005;2(4):281–293. doi:10.1098/rsif.2005.0042.

78. Centers for Disease Control Taiwan. Taiwan National Infectious Disease Statistics System;. Available from: http://nidss.cdc.gov.tw/en/SingleDisease.aspx?dc=1{&}dt=4{&}disease=061{&}position=1.

79. Chang SF, Huang JH, Shu PY. Characteristics of dengue epidemics in Taiwan. J Formos Med Assoc. 2012;111(6):297–299. doi:10.1016/j.jfma.2011.12.001.

80. Cobelli C, DiStefano JJ. Parameter and structural identifiability concepts and ambiguities: a critical review and analysis. Am J Physiol. 1980;239(1):R7–R24.

81. Audoly S, Bellu G, D’Angi`o L, Saccomani MP, Cobelli C. Global identifiability of nonlinear models of biological systems. IEEE Trans Biomed Eng. 2001;48(1):55–65. doi:10.1109/10.900248.

82. Raue A, Kreutz C, Maiwald T, Bachmann J, Schilling M, Klingm??ller U, et al. Structural and practical identifiability analysis of partially observed dynamical models by exploiting the profile likelihood. Bioinformatics. 2009;25(15):1923–1929. doi:10.1093/bioinformatics/btp358.

83. Miao H, Xia X, Perelson AS, Wu H. On identifiability of nonlinear ODE models and applications in viral dynamics. SIAM Rev. 2011;53(1):3–39. doi:10.1137/090757009.

84. Ollivier F. Le probleme de l’identifiabilite structurelle globale: approche theorique, methodes effectives et bornes de complexite. École Polytechnique; 1990.

85. Pia Saccomani M, Audoly S, Bellu G, D’Angio L. A new differential algebra algorithm to test identifiability of nonlinear systems with given initial conditions. In: Proc. 40th IEEE Conf. Decis. Control (Cat. No.01CH37228). IEEE;. p. 3108–3113. Available from: http://ieeexplore.ieee.org/lpdocs/epic03/wrapper.htm?arnumber=980295.

86. Meshkat N, Anderson C, DiStefano III JJ. Alternative to Ritt’s pseudodivision for finding the input-output equations of multi-output models. Math Biosci. 2012;239(1):117–123. doi:10.1016/j.mbs.2012.04.008.

87. Eisenberg M. Generalizing the differential algebra approach to input-output equations in structural identifiability; 2013. Available from: http://arxiv.org/abs/1302.5484.

88. Rothenberg TJ. Identification in Parametric Models. Econom J Econom Soc. 1971;39(3):577–591.

89. Cintrón-Arias A, Banks HT, Capaldi A, Lloyd AL. A sensitivity matrix based methodology for inverse problem formulation. J Inverse Ill-posed Probl. 2009;17(6):545–564. doi:10.1515/JIIP.2009.034.

90. Eisenberg MC, Hayashi MAL. Determining identifiable parameter combinations using subset profiling. Math Biosci. 2014;256:116–126. doi:10.1016/j.mbs.2014.08.008.

91. Jacquez JA, Greif P. Numerical parameter identifiability and estimability: Integrating identifiability, estimability, and optimal sampling design. Math Biosci. 1985;77(1-2):201–227. doi:10.1016/0025-5564(85)90098-7.

92. Meshkat N, Kuo CEz, DiStefano J. On Finding and Using Identifiable Parameter Combinations in Nonlinear Dynamic Systems Biology Models and COMBOS: A Novel Web Implementation. PLoS One. 2014;9(10):e110261. doi:10.1371/journal.pone.0110261.

93. Bellu G, Saccomani MP, Audoly S, D’Angiò L. DAISY: A new software tool to test global identifiability of biological and physiological systems. Comput Methods Programs Biomed. 2007;88(1):52–61. doi:10.1016/j.cmpb.2007.07.002.

94. Wu HH, Wang CY, Teng HJ, Lin C, Lu LC, Jian SW, et al. A Dengue Vector Surveillance by Human Population-Stratified Ovitrap Survey for Aedes (Diptera: Culicidae) Adult and Egg Collections in High Dengue-Risk Areas of Taiwan. J Med Entomol. 2013;50(2):261–269. doi:10.1603/ME11263.

95. Morrison AC, Zielinski-Gutierrez E, Scott TW, Rosenberg R. Defining Challenges and Proposing Solutions for Control of the Virus Vector Aedes aegypti. PLoS Med. 2008;5(3):e68. doi:10.1371/journal.pmed.0050068.

96. Chang MS, Christophel EM, Gopinath D, Abdur RM. Challenges and future perspective for dengue vector control in the Western Pacific Region. West Pacific Surveill Response. 2011;2(2):e1–e1. doi:10.5365/wpsar.2010.1.1.012.

97. Benelli G, Jeffries C, Walker T. Biological Control of Mosquito Vectors: Past, Present, and Future. Insects. 2016;7(4):52. doi:10.3390/insects7040052.

